# Scale-free dynamics of cerebrospinal fluid regions is associated with Alzheimer’s disease-related pathology

**DOI:** 10.1101/2024.04.21.589866

**Authors:** Qiwei Guo, Lin Hua, Zhiying Zhao, Zhen Yuan

**Author notes:** These authors contributed equally to this work.

## Abstract

Cerebrospinal Fluid (CSF) clearance is one of the central mechanisms for removal of amyloid-β (Aβ) and tau proteins from the central nervous system that are closely linked to Alzheimer’s Disease (AD) pathology. Conventional functional imaging studies in AD have primarily focused on activities in the brain while ignoring macroscopic cerebrospinal fluid (CSF) activities. In the current study, we utilized a public dataset from the Alzheimer’s Disease Neuroimaging Initiative. Our analysis spanned both brain and cerebrospinal fluid (CSF) areas. We compared the scale-free dynamics in fMRI signals across three groups: cognitively normal individuals (CN), people with Alzheimer’s Disease (AD), and those with mild cognitive impairments (MCI). Scale-free dynamics are patterns of brain activity that remain consistent across different time scales. Our comparison focused on the Hurst exponent (H), a measure derived from detrended fluctuation analysis (DFA), to characterize these dynamics. This comparison revealed clusters in the fourth ventricle and subarachnoid space (medioventral channel along the spinal axis). In these CSF-filled structures, H was significantly higher in AD group comparing to CN. Moreover, H in these two clusters correlated with AD pathological biomarkers including CSF Aβ and tau in the AD group. These findings suggested scale-free properties of macroscopic CSF flow as a potential imaging biomarker for AD. This biomarker can be readily acquired from common resting-state fMRI scans and therefore may be valuable for AD diagnosis.

## Introduction

Alzheimer’s Disease (AD) is one of the most common types of dementia in elder adults representing a major public health concern that brings serious social and economic burdens. Acute decline of memory function is one of the early symptoms of AD. With the development of the disease, patients develop severe memory impairment, loss of daily life and self-care ability, and other cognitive and emotional disability symptoms and behavioral abnormalities (1). The etiology of Alzheimer’s disease (AD) remains elusive, and a definitive pharmacological intervention for its cure has yet to be established (2). The pathogenesis of AD is believed to be caused by the accumulation of toxic waste such as Amyloid β (Aβ) protein and tau protein, and this accumulation is recently linked with impairments in glymphatic system which removes these waste proteins from the brain The glymphatic system, a critical waste clearance pathway in the brain, operates intricately with cerebrospinal fluid (CSF) dynamics. This interplay is pivotal for maintaining brain health and offers a unique lens through which we can examine the underlying mechanisms of neurological disorders. (3–5).

Brain Functional magnetic resonance imaging (fMRI) represents a useful non-invasive approach to investigate the alterations in brain functions in AD patients. Evidence from resting-state fMRI converges on the abnormal connectivity in default mode network, salience network and limbic system (6). Overall, fMRI technology has made a great stride in biomarker research for AD over the past decade (7).

A wide range of biomedical signals exhibit recurring spatial or temporal dynamics that appeared to be “scale-free” including the fMRI signal (8,9).This term refers to a type of behavior in a system where the same patterns occur at many different scales or sizes. In the CSF dynamics, it means that the patterns of how the fluid moves or changes can be similar whether you’re looking at small or large volumes or over short or long time periods. The structural characteristics of signals are often visually apparent, but not captured by conventional measures like the average amplitude of the signal. Understanding these scale-free dynamics in biomedical signals, such as fMRI, is crucial for deciphering complex biological processes. To this end, fractal analysis, particularly through the Hurst exponent (H) derived from detrended fluctuation analysis (DFA), offers a nuanced approach for examining these patterns in the context of brain fMRI. The Hurst exponent (H), a key metric derived from detrended fluctuation analysis (DFA), helps us understand the memory and predictability of a system. When H is between 0 and 1, it suggests that the system’s future behavior is less influenced by its past behavior, akin to a more random or unpredictable process. In contrast, H values between 1 and 2 indicate that the system’s future behavior is more strongly influenced by its past, showing a higher degree of predictability or memory. In the context of brain fMRI, these H values help us discern different types of dynamics within brain tissue and functional networks (10).

It has been reported that H varies between different brain tissue types and functional networks in many fMRI studies(8,9). Increased cognitive effort leads to a decrease in H, reflecting changes in brain’s scale-free signal structuring in response to task complexity and novelty (11). Aging and progression of neurodegenerative disease were found to be associated with scale-invariant properties in brain dynamics, as evidenced by a positive correlation between age and average H in brain gray matter, particularly in parietal and frontal lobes (12) Variations in H in the grey matter may help in identifying early signs of neurological disease, assessing disease severity and monitoring treatment effects (12,13). These findings have highlighted H as a useful index for accessing cognitive function-related changes in the brain.

Traditionally, in both resting-state and task-based fMRI studies, including those focusing on neurodegenerative diseases, macroscopic CSF activity has often been dismissed as a nuisance signal that interferes with the BOLD signal. Consequently, it is typically removed during the data preprocessing stage, usually through regression techniques This approach to data ‘denoising’ may neglect significant alterations in CSF dynamics that could signal glymphatic clearance issues in Alzheimer’s Disease (AD). Highlighting the value of reevaluating CSF dynamics, recent groundbreaking research by Fultz et al. revealed that CSF inflow in the ventricles is driven by widespread cortical hemodynamic oscillation (14). Following this unique path, Han et al. has established a link between neuro-fluid coupling and disease markers in AD and Parkinson’s Disease (PD), suggesting that fMRI can uncover CSF activity changes with pathological significance (15,16). These findings suggested that fMRI can detect changes in CSF activity that were of pathological importance. Building on this insight and considering that the Hurst exponent (H) provides unique information on BOLD activity and fluid dynamics (17), our exploratory study aims to investigate H’s potential to identify AD-related scale-free changes in both brain and CSF activities using data from the Alzheimer’s Disease Neuroimaging Initiative (ADNI).

## Methods

### Participants

We included resting state fMRI, formal diagnosis, and CSF biospecimen data in 287 participants from the Alzheimer’s Disease Neuroimaging Initiative (ADNI) database initially. The ADNI study received ethical approval from the institutional review boards of all participating centers, and written informed consent was obtained from all participants or their authorized representatives. Participants who had both structural and resting-state functional images collected on 3 Tesla MRI systems were included. After data quality screening, one participant was excluded due to excessive head motions that over 3 mm threshold, resulting in a total of 286 participants who entered the formal data analyses. The demographic information of both groups is reported in Table 1.

**Table 1.** Participants characteristics.

| Characteristics | AD(n=83) | MCI(n=123) | NC(n=80) | p-value (statistics of tests) |
| --- | --- | --- | --- | --- |
| Age (years), mean(s.d.) | 74.51 (7.48) | 72.86 (7.86) | 71.84 (6.50) | 0.058($F_{2,283} = 2.87$) |
| Sex, n (% of female) | 33 (40.96%) | 60 (48.78%) | 50 (62.50%) | 0.01 ($\chi^2=8.56$) |
| Education (years), mean (s.d.) | 15.57 (2.64) | 16.26 (2.57) | 15.98 (2.40) | 0.22 ($F_{2,250} = 1.54$) |
| MMSE, mean<br>(s.d) | 22.76 (3.05) | 27.52 (1.86) | 28.71 (1.37) | < .001 ($F_{2,282} = 181.06$) |
| CSF Aβ42,<br>mean(s.d.) | 339.65 (283.75) | 971.63 (623.31) | 930.26 (760.34) | < .001 ($F_{2,153} = 17.49$) |
| CSF pTau<br>mean(s.d.) | 46.71 (23.28) | 29.45 (18.92) | 21.90 (9.77) | < .001 ($F_{2,153} = 24.92$) |
| CSF Tau, mean<br>(s.d.) | 223.82(149.94) | 282.84(154.09) | 194.91(106.95) | 0.005 ($F_{2,153} = 5.58$) |
| pTau/Aβ42 ratio | 0.278 (0.288) | 0.047 (0.054) | 0.053 (0.066) | < .001 ($F_{2,153} = 30.67$) |
| APOE ε4 status<br>(% of carriers) | 44 (53.01%) | 36 (29.27%) | 25 (31.25%) | < .001 ($F_{2,218} = 16.20$) |

Abbreviations: MMSE, mini-mental state examination; CSF, cerebrospinal fluid; APOE, Apolipoprotein E; s.d., standard deviation; Tau, Tau protein level; pTau, hyperphosphorylated tau protein level. Aβ42, amyloid beta peptide 42 level.

### Resting-state fMRI data preprocessing

Imaging data preprocessing was performed using CONN toolbox (release 21.a, https://web.conn-toolbox.org/). Resting-state data from all participants underwent an identical preprocessing pipeline in the order of realignment with correction of susceptibility distortion interactions, slice timing correction, outlier detection, direct segmentation and MNI-space normalization, and spatial smoothing. These were first realigned using realign & unwarp procedure based on SPM, where all scans were co-registered to a reference image (first scan of the series) using a least squares approach and a 6-parameter (rigid body) transformation and resampled using b-spline interpolation to correct for motion and magnetic susceptibility interactions. Temporal misalignment between different slices of the functional data was corrected following SPM slice-timing correction (STC) procedure, using sinc temporal interpolation to resample each slice to a common mid-acquisition time. Potential outlier scans were identified by ART (18) as acquisitions with framewise displacement above 0.9 mm or global BOLD signal changes above 5 standard deviations, and a reference BOLD image was computed for each subject by averaging all scans excluding outliers. Functional and anatomical data were normalized into standard MNI space, segmented into grey matter, white matter, and CSF tissue classes, and resampled to 3 mm^3^ isotropic voxels following a direct normalization procedure using SPM unified segmentation and normalization algorithm with the default IXI-549 tissue probability map template. Last, functional data were smoothed with a Gaussian kernel of 6 mm full-width half-maximum (FWHM). Given the of evidence, nuisance regression of ROI (average global BOLD, CSF, white matter) is omitted as well as the bandpass filtering this study as we have interest in CSF and gBOLD signal and had no hypothesis on the frequency range of CSF dynamics in relation to pathological aging.

### Detrended fluctuation analysis (DFA)

DFA has been proven to outperform other approach in similar situations (19–22).The Hurst exponent (H) estimation by DFA method has become a frequently used tool for time domain fractal analysis allowing for its application in estimating the fractal property of short time series. The Hurst exponent was estimated by Detrended Fluctuation Analysis (DFA) which briefly consists of following steps and can be further described at (23):

1. Converting noise-like time series into a random walk time series by taking the cumulative sum of the difference between the overall mean and value of each time point.
2. Choosing a window size, then dividing the whole time series into same length, non-overlapping segments.
3. Estimating local fluctuation of each segment by taking the root mean square (RMS) of residual variation of each time point. The residual variation is computed through the subtraction between fitted value and y value of each time point using ordinary least square (OLS) method. A range of different window size is necessary to capture different temporal fluctuations. Small window sizes capture fast fluctuation whereas large windows capture slow fluctuation. Here we used a window scale vector of [5, 10, 15, 20, 25, 30, 35] for our estimation, a configuration that has been previously used for fMRI data. Results derived from different windows size series are reported in Supplement.
4. The overall fluctuation (overall RMS) can be summarized by all segments of each window size by taking the RMS of previous local RMSs.
5. Each selection of time window yields a corresponding overall fluctuation, making it the overall RMS as a function of window scales. Then Hurst exponent is calculated from the linear slope of this function.

### Among-groups comparison of Hurst exponent

Hurst exponent (H) of each voxel was estimated from the preprocessed resting-state functional images to generate whole brain maps using DFA method described above. Non-brain voxels were removed by applying an inclusive brain mask provided by DPABI 7.0 which also covers the CSF areas outside the brain (Fig S2 – overlay the brain mask and segmented group-mean CSF on MNI brain with some transparency). Analysis of variance (ANOVA) was performed in SPM12 (release 12.7771, https://www.fil.ion.ucl.ac.uk/spm/software/spm12/) to identify regional differences in H maps between AD, MCI and CN groups at a whole brain level. Clusters remained significant after correcting for Family-Wise Errors (FWE) (corrected P < 0.001, k>30) was selected as regions of interest (ROI) for exploratory analyses by extracting the average H values from the ROIs.

## Results

### Participants characteristics

Table 1 has summarized demographical characteristics of 286 participants with statistics. The imaging and biospecimen data were available in 83 participants in AD (Alzheimer’s Disease) group, 80 participants in CN (Normal control) group and 123 participants in MCI (Minor Cognitive Impairment) group. There was a significant difference in sex ratio among participant groups as well as the mini-mental state examination score (MMSE), CSF Amyloid Beta peptide 42 level (CSF Aβ42), CSF ptau-A42 ratio (P-tau/Aβ42 ratio), CSF Tau protein level (CSF Tau), CSF partial phosphorylated Tau level (CSF pTau).

CSF clusters showing differences in Hurst exponent (H) between AD, MCI and NC groups. Voxel-wise H maps that calculated from DFA are shown for each group are shown (Fig S1). H demonstrated a significant decreasing trend in Grey Matter (GM) in participants with MCI and AD comparing to CN (Mean GM-H of AD= 0.919, SD = 0.232; Mean GM-H of MCI = 0.824, SD = 0.283; Mean GM-H of CN = 0.728, SD = 0.195; ANOVA F = 11.32, P < 0.001; See Fig S2a for post hoc tests.). A spatial analysis of variance (ANOVA) has been applied to three groups to find the spatial difference of H while adjusting for age, sex, and education using SPM. The result revealed two clusters showing location of significant differences among three groups (Fig 1). One in the fourth ventricle (peak F_2,281_ = 21.18, 44 voxels) and a cluster in medioventral subarachnoid space near the spinal cord above craniocervical junction (peak F_2,281 =_ 24.83, 59 voxels) after FWE correction. Treating them as a Region of Interest (ROI), mean H of these two clusters was extracted for each participant and referred to as CSF-H hereinafter.

**Fig 1.**
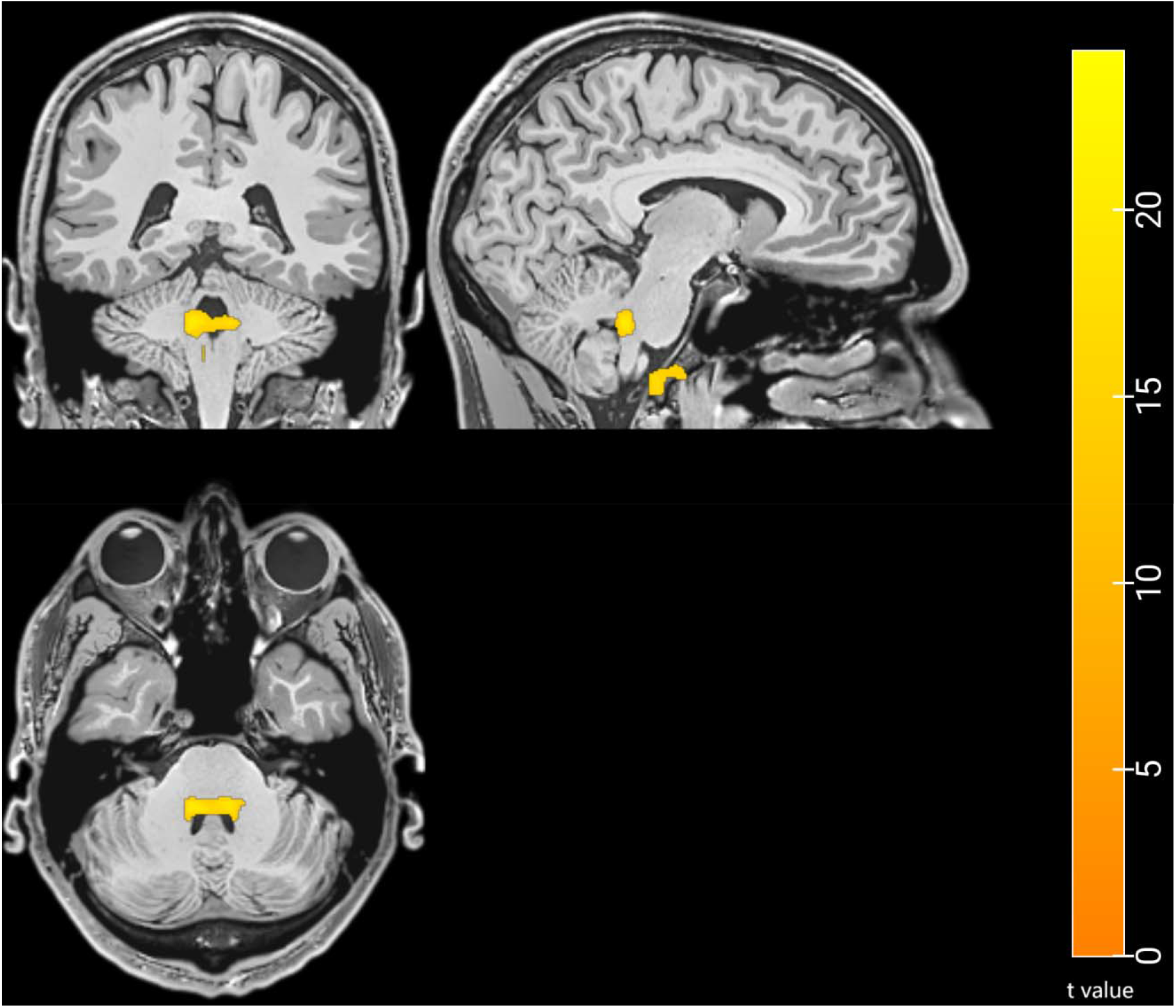
Differential Clusters for averaged Hurst maps among AD, MCI, and NC groups by ANOVA. Clusters showing difference in H between AD, MCI and CN groups in whole-brain ANOVA adjusted for age and sex. The clusters survived from a height threshold F = 14.90 with an extent threshold k = 30 voxels. Color bar corresponds to t value.

### Relationship between CSF-H and AD risk factors

We then investigated the relationship of CSF-H with demographic characteristics including age, sex, education, and status of Apolipoprotein E (APOE) genotype. We first observed a significant decreasing trend among AD, MCI, and CN groups (Mean CSF-H of AD= 0.874, SD = 0.312; Mean CSF-H of MCI = 0.643, SD = 0.232; Mean CSF-H of CN = 0.561, SD = 0.236; ANOVA F = 32.86, P < 0.001; See Fig 2a for post hoc tests.). We did not observe significant correlation between CSF-H and Age (Pearson’s R = 0.094, P = 0.11; Fig 2a.) or between CSF-H and Sex (Student’s t = 0.734, P = 0.46, n = 286; Fig 2b.). We found a weak correlation between H and education only in MCI group (Pearson’s R = -0.22, P = 0.029; Fig 2c.). We then compared CSF-H between phenotypes while controlling for age and sex. Consistent with our whole-brain ANOVA finding, CSF-H differed significantly between the groups (ANOVA F_2,283_ = 32.86, P < 0.001; Fig 2d) with the CN showing lower CSF-H than the MCI (mean difference: -0.082, t = -2.198, P_tukey_ = 0.073) and AD groups (mean difference: 0.313, t = 7.71, P_tukey_ < 0.001). AD group also showed higher CSF-H comparing to the MCI group (mean difference: 0.231, t =6.281, P_tukey_ < 0.001). CSF-H was also compared across different apolipoprotein E (APOE) gene statuses and showed a significantly dose-dependent relationship with the copies of APOE ε4 alleles (ANOVA F_2,218_ = 3.44, P = 0.034, Fig 2e.).

**Fig 2.**
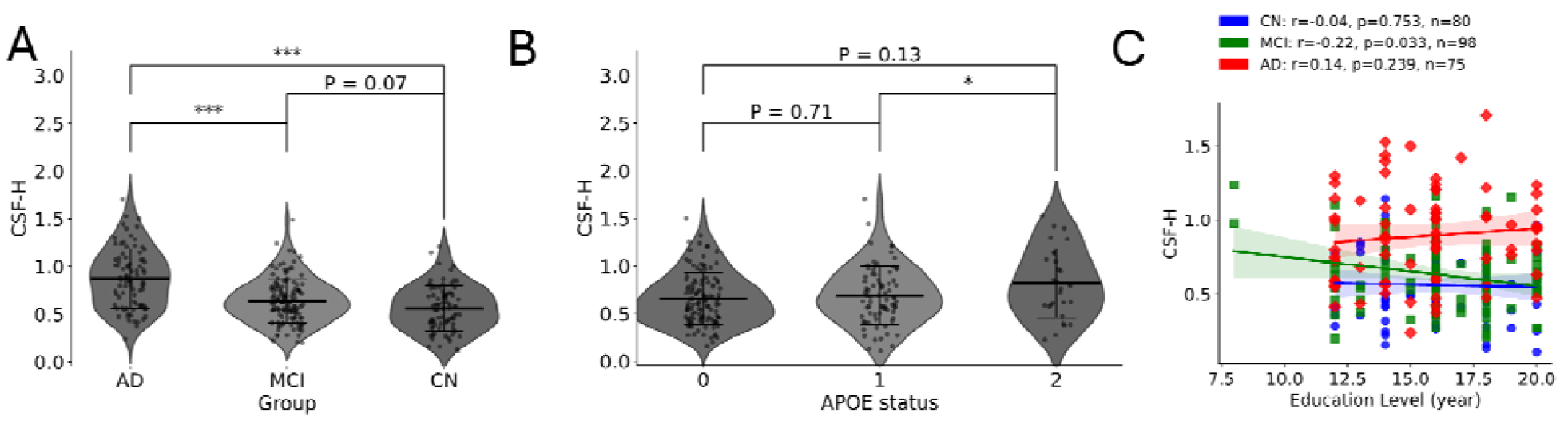
Relationship between CSF-H and AD risk factors. (a) Comparison of Average CSF-H among AD, MCI and CN group visualized in violin plot. The upper and lower whiskers mark full range of distribution excluding outliers while the line in the middle denotes median. Middle line, the mean value of H. Tuckey correction was used for post hoc tests. Error bar represents mean ± SEM; *** P < 0.001, ** P < 0.01, *P < 0.05, ns. not significant, same notion was used for next panel. (b) Average H value in different APOE status groups. A Violin plot was made to analyze the average CSF-H in different APOE status groups. The upper and lower cutoff stands for standard error. Middle line represents the mean value of H. APOE status 0 represents one carries zero apolipoprotein E ε4 allele while 1 means subject carries one APOE ε4 allele and 2 means a carrier with 2 alleles. (c) Group-wise correlation analysis, education vs. CSF-H. A Scatter plot was made to view the relationship between CSF-H and a risk factor, education. Solid line with corresponding colors (red, green, and blue) and symbols represents the fitted linear relationship within each group while black line denotes this relationship in all the participants. Shadow areas cover the 95% confidence interval (CI) with respect to each group. The color scheme is maintained consistent with previous figures and Pearson’s correlation coefficient is used for the rest of this figure.

### Relationship between CSF-H and AD-related pathology

CSF-H was then correlated with symptom indicators of AD, namely MMSE score, CSF pTau and CSF Aβ42 levels. After adjusting for age and sex, a significant negative correlation (Pearson’s r = -0.378, P *<* 0.001, n=285) was found between CSF-H and MMSE score across all condition groups. However, this linear relationship was not significant when examined separately for each group (Fig 3a). We found an overall significant positive correlation (Pearson’s r =0.36, P <0.001, n=156) between CSF-H and CSF pTau level in the entire sample. Specifically, we found the correlation was mainly driven by the AD group (Pearson’s r = 0.40, P = 0.005, n = 49; Fig 3b) but not the other groups. Regarding the relationship between CSF-H and CSF Aβ42 level, we found a significant negative correlation across all groups (Pearson’s r = -0.21, P = 0.009, n = 156). Specifically, we found this overall correlation was also mainly driven by the AD group (Pearson’s r = -0.50, P < 0.001, n=49) but no other groups (Fig 3c).

**Fig 3.**
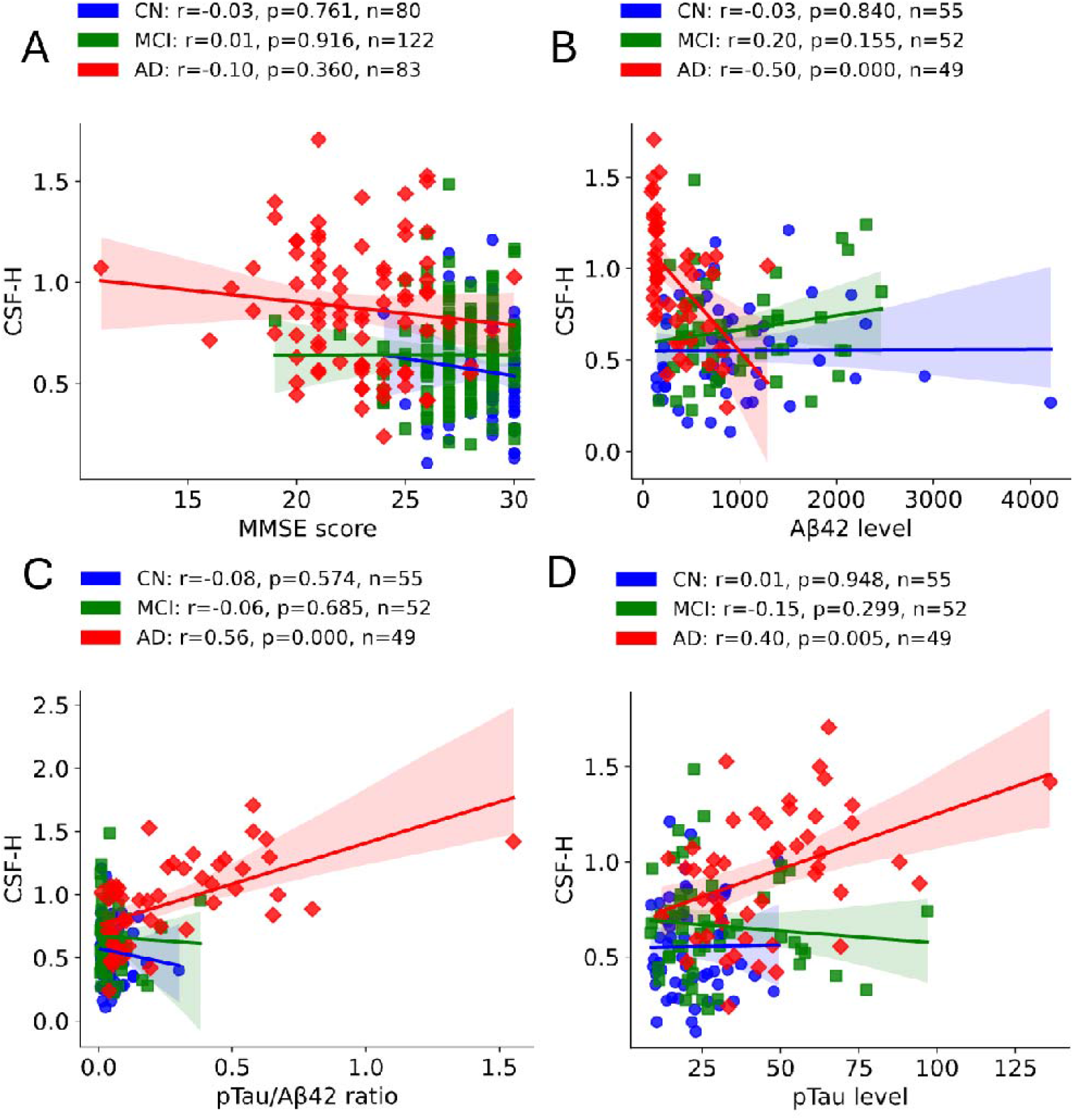
CSF-H is correlated with AD pathological markers. (a) Group-wise correlation analysis, education vs. CSF-H. A scatter plot was made to visualize and estimate the dependence of CSF-H on MMSE score. Solid line with corresponding colors (red, green and blue) and symbols represents the fitted linear relationship within each group while black line denotes this relationship in all the participants. Shadow areas cover the 95% confidence interval. The color scheme is maintained consistent with previous figures and Pearson’s correlation coefficient is used for the rest of this figure. is used throughout the rest of the manuscript. (b) Group-wise correlation analysis, CSF pTau vs. CSF-H. A scatter plot was made to view the relationship between CSF-H and a pathological biomarker, CSF pTau level. (c) Group-wise correlation analysis, CSF Aβ42 vs. CSF-H. A scatter plot was made to view the relationship between CSF-H and a pathological biomarker, CSF Aβ42 level. Group-wise correlation analysis, CSF pTau/Aβ42 ratio vs. CSF-H. A scatter plot was made to view the relationship between CSF-H and pTau/Aβ42 ratio.

CSF pTau/Aβ42 ratio was suggested as a promising predictor for AD pathological status including amyloid PET status and cognitive decline. We therefore further explored whether this biomarker was linked to CSF-H in our sample. Indeed, ANOVA suggested that pTau/Aβ42 ratio also differed between the groups (ANOVA F_2,153_ = 30.67, P < 0.001). Not only CSF-H correlated with pTau/Aβ42 across all participants (Pearson’s r = 0.50, P < 0.001, n = 156) but also this correlation was driven by the trend in AD group (Pearson’s r = 0.56, P < 0.001, n = 49; Fig 3d).

## Discussion

In this study, we performed fractal analysis of resting state fMRI signals in AD, MCI and heathy controls using a data-driven approach. Comparison of the scale-free property between the three group of participants revealed two regions in CSF space, namely the fourth ventricle and subarachnoid space (SAS) in which Hurst exponent was higher in AD group comparing to MCI and HC. Moreover, CSF-H correlated with cognitive function (MMSE) and pathological biomarkers including P-tau, T-tau, and Aβ42 in CSF in AD but no other two groups, suggesting that this alteration in CSF-H is disease-specific.

The pathogenesis of AD has been attributed to the accumulation of toxic protein aggregates (24) which can be accelerated by deficient clearance of these waste products. The extracellular removal of Aβ and tau proteins relies on CSF transport(25) in various pathways. CSF was primarily produced in choroid plexuses in the ventricles of the brain. CSF then flows to subarachnoid space (SAS) through the median aperture. A portion of CSF in SAS continues to circulate into brain parenchyma through peri-artery space and exchange with interstitial fluid (ISF) via aquaporin-4 (AQP4) channels. This circulation bathes and metabolizes the toxic waste of the brain(26). Therefore, our findings in ventricular and SAS CSF dynamics may reflect CSF clearance efficiency which is weakened in AD patients(27). Vasomotion and cardiac pulsations are the main drivers for CSF oscillations in resting-state fMRI, both of which are influenced by aging and aging-related diseases. The association between CSF-H and waste protein load in CSF showed that this index quantifies clearance functioning, although more studies are needed to reveal the underlying mechanisms.

Despite the crucial role of CSF circulation in AD pathogenesis, functional imaging studies have rarely examined whether dysregulated CSF flow is involved in AD. Studies using phase-contrast MRI showed that CSF pulsations are heterogeneous in terms of temporal dynamics and directionalities (28–30) across different compartments, which are challenging to characterize using a single index that is pathologically meaningful for neurodegenerative diseases. The cardiac and vascular contributions to CSF pulsations are not included in the typical 0.001-0.01 Hz band-pass filtering range in conventional resting-state fMRI studies (30). Here we showed that omitting filtering in turn preserves CSF dynamics that are linked to AD pathology which can be quantified using Hurst exponent. In our study, we conducted an unbiased whole-brain estimation of the Hurst exponent (H) across three groups to explore any potential differences. Interestingly, this approach led us to identify notable changes in CSF dynamics, a finding that emerged without a prior hypothesis and aligns with recent research in the field (14–16). This serendipitous discovery underscores the value of unbiased, exploratory analyses in uncovering new aspects of disease pathology, particularly in the context of Alzheimer’s Disease. However, this also warrants further examinations on the link between CSF circulation outside of brain and extracellular clearance in the brain through CSF-ISF exchange. Moreover, it’s of great importance to learn what leads to dysregulated CSF flow in AD as well as the CSF circulation between sleep and awake states.

In summary, we reported evidence that the mono-fractal property of ventricular and subarachnoid CSF oscillations discriminates AD patients from MCI and CN. This measurement associated with cognitive function across groups, notably, its correlation with CSF Aβ42 and tau levels was only present in AD group. These findings provided unique angles for understanding impairments in CSF clearance in neurodegenerative diseases and suggested the potential of CSF-H as a novel and in-expensive marker for AD diagnosis. Replications and further investigations are needed to validate the current findings.

## Acknowledgement

This work was supported by Macao Science and Technology Development Fund to study the role of macroscopic CSF flow in brain functions and development (FDCT 0015/2023/ITP1 to ZZ)

## Supporting information

**S1 Fig.**
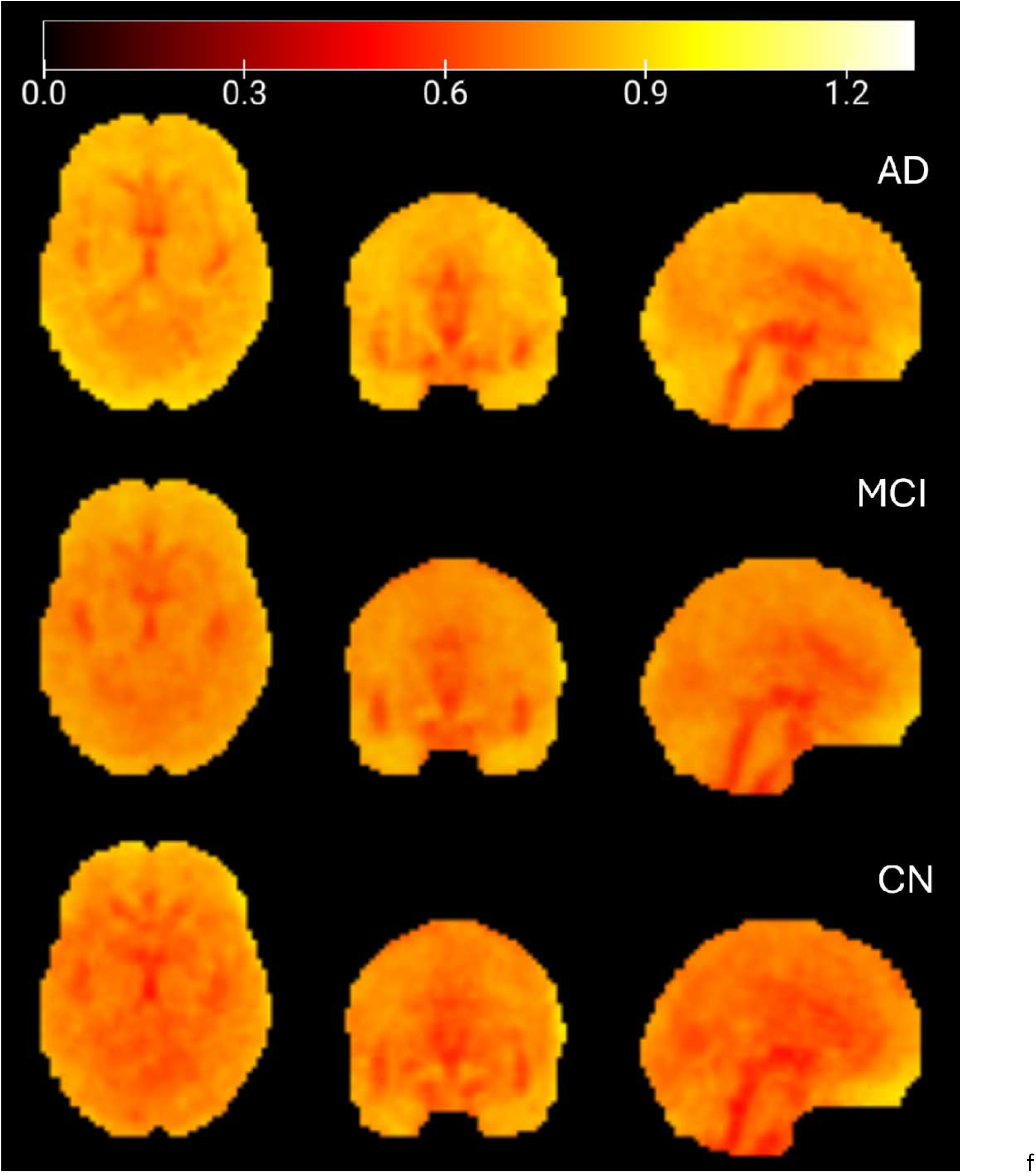
Group averaged Hurst exponent (H) maps for AD, MCI, and CN groups. Group labels indicates group names, AD = Alzheimer’s Disease; MCI = Mild Cognitive Impairment; CN = Cognitively Normal. Color bar corresponds to H value.

**S2 Fig.**
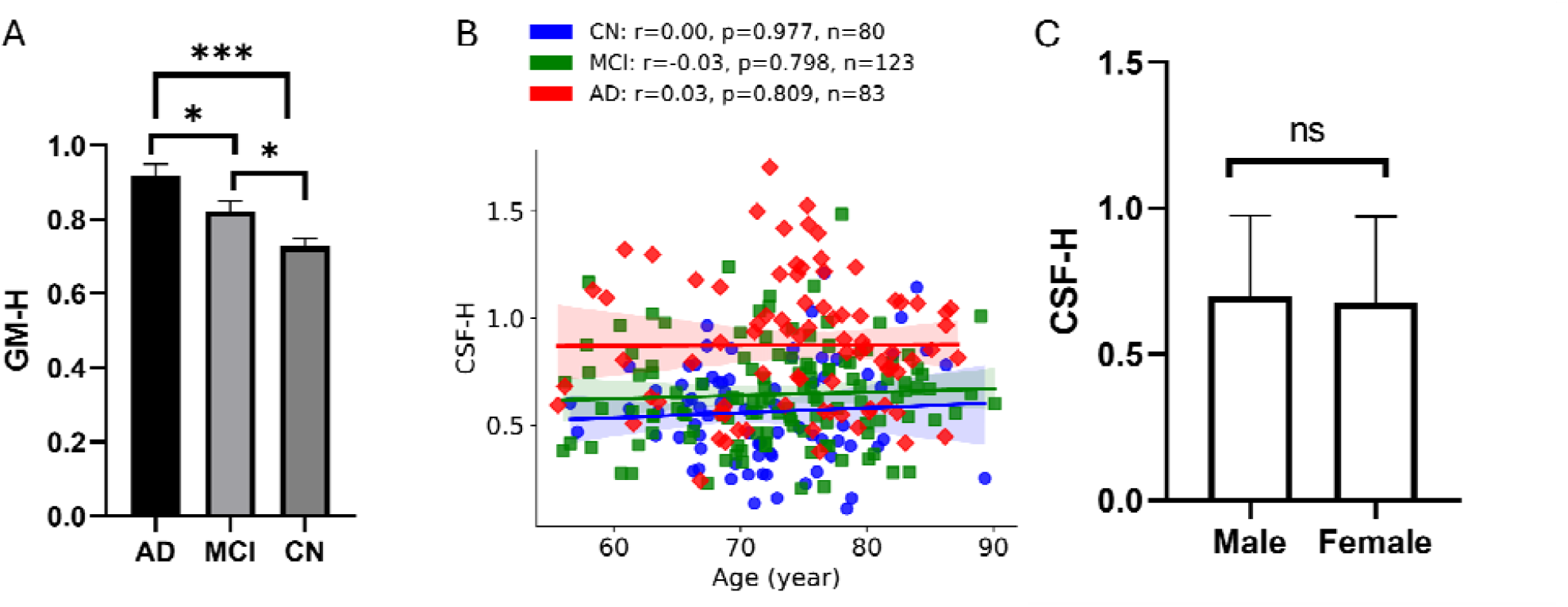
Relationship between CSF-H and AD risk factors. a) Comparison of average GM-H among AD, MCI and CN group. Tuckey correction was used for post hoc tests. Error bar represents mean ± SEM; *** P < 0.001, ** P < 0.01, *P < 0.05, ns. not significant. b) Group-wise correlation analysis, Age vs. CSF-H. A scatter plot showing the relationship between CSF-H and age. Solid line with corresponding colors (red, green and blue) and symbols represents the fitted linear relationship within each group while black line denotes this relationship in all the participants. Shadow areas cover the 95% confidence interval. c) Average CSF-H for male and female. Error bar stands for standard error. The ns label is the abbreviation of not significant.

**S3 Fig.**
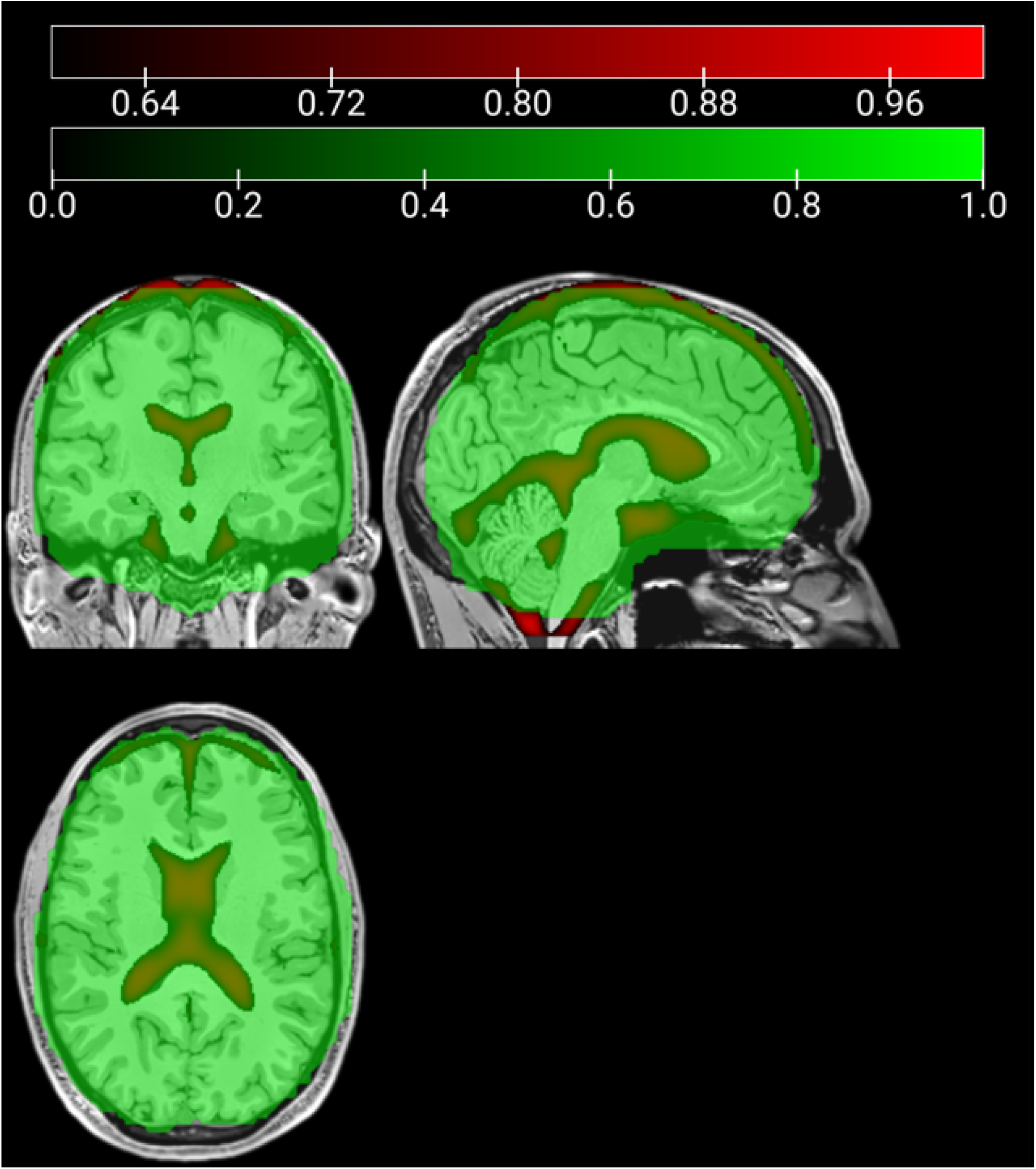
brain region coverage in this analysis. The group analysis is covered in red region comparing to standard MNI template. Red spectrum represents an overlay of group average image of CSF segmentation. Green spectrum represents the brain mask used in this study.

**S4 Fig.**
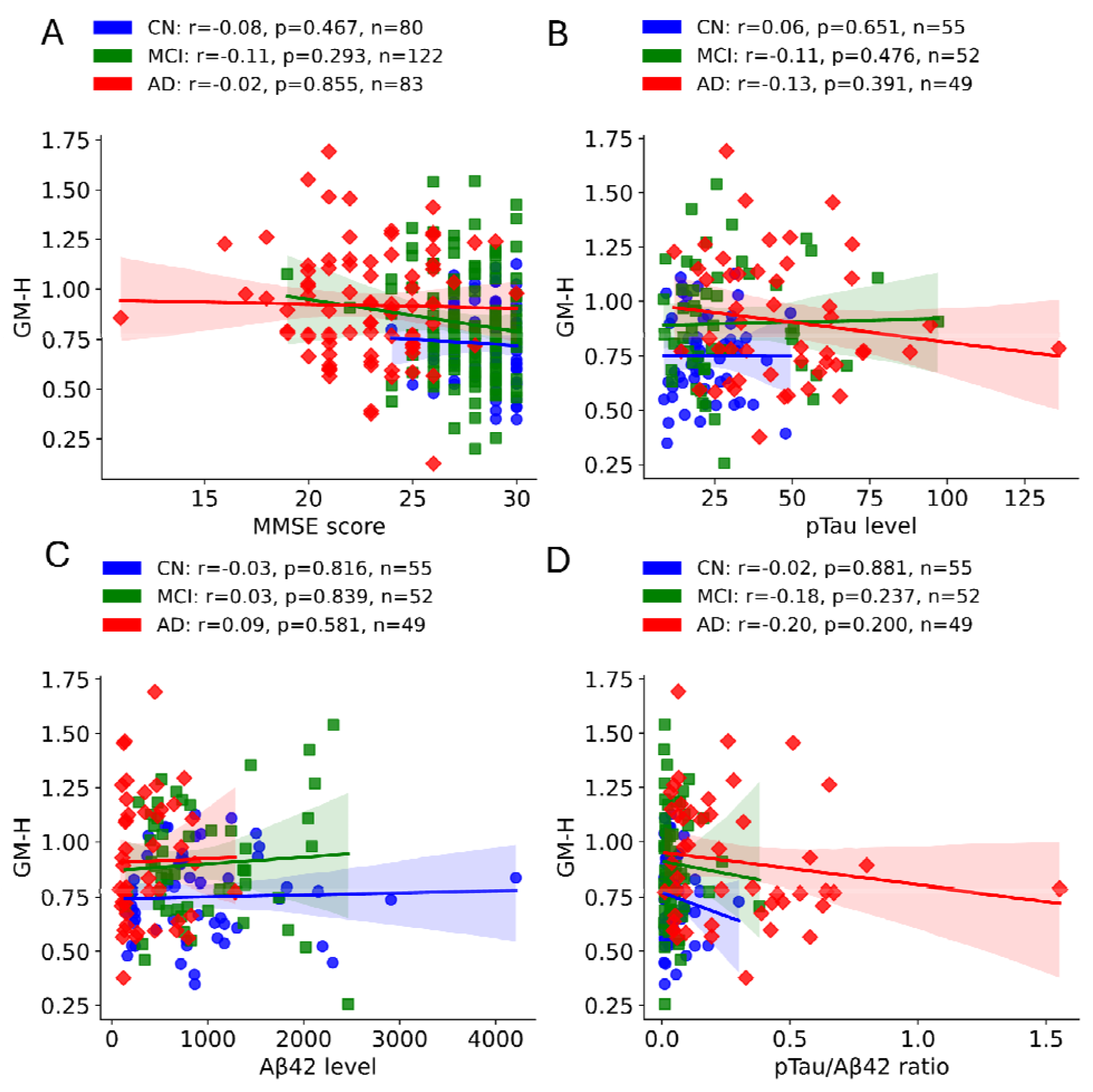
GM-H is not significantly correlated to any of pathological markers. (a) Group-wise correlation analysis for education vs. GM-H. A scatter plot was made to visualize and estimate the dependence of GM-H on MMSE score. Solid line with corresponding colors (red, green and blue) and symbols represents the fitted linear relationship within each group while black line denotes this relationship in all the participants. Shadow areas cover the 95% confidence interval. This color scheme is used throughout the rest of the manuscript. The color scheme is maintained consistent with previous figures and Pearson’s correlation coefficient is used for the rest of this figure. (b) Group-wise correlation analysis, CSF pTau vs. GM-H. A scatter plot was made to view the relationship between GM-H and a pathological biomarker, CSF pTau level. (c) Group-wise correlation analysis, CSF Aβ42 vs. GM-H. A scatter plot was made to view the relationship between GM-H and a pathological biomarker, CSF Aβ42 level. Group-wise correlation analysis, CSF pTau/Aβ42 ratio vs. GM-H. A scatter plot was made to view the relationship between GM-H and pTau/Aβ42 ratio.

